# Mechanical Induced Stress Enhances Arabidopsis Susceptibility to *Fusarium oxysporum* in a Jasmonic Acid-Dependent Manner

**DOI:** 10.64898/2026.11.24.769068

**Authors:** Eric Brenya, Mahfuza Pervin, Avijit Biswas, Lucky Paudel, Zhong-Hua Chen, David T Tissue, Scott Johnson, Christopher Ian Cazzonelli

## Abstract

Prolonged mechanical-induced stress (MIS) primes juvenile Arabidopsis plants for enhanced resistance against necrotrophic pathogens through the activation of jasmonate-mediated defense pathways. We hypothesised that MIS would weaken defense against hemibiotrophic pathogens that exploit jasmonate pathways to promote infection. We show that touch-acclimated wild-type plants were more susceptible to *Fusarium oxysporum* f. sp. *conglutinans* (Foc) infection than un-touched control plants. Jasmonic acid (JA) levels as well as JA defense-related gene expression were hyper-induced for up to 5 days post-inoculation in the MIS plants. In contrast, the MIS- and control-JA biosynthesis-deficient mutant *jasmonate resistance 1* (*jar1-11*) was more resistant to Foc infection compared to the WT. The *jar1-11* mutant showed significant attenuation in the induction of JA defense-responsive gene expression. In the absence of Foc infection, a single event of MIS similarly induced JA biosynthesis and stress-associated marker genes, but not JA signalling genes, in naïve and touch-acclimated plant tissues. Touch-acclimated plants had higher basal levels of JA and JA defense-related genes expression, including *PDF1.2*, *VSP1,* and *LOX1*. Our findings highlight that prolonged MIS of Arabidopsis cross-primes JA-dependent susceptibility to a hemibiotrophic pathogen that can hijack the JA pathway to promote infection, revealing a potential trade-off when utilising MIS to promote immunity against necrotrophic pathogens or alter morphological traits for agronomical gains.

**Highlights:**

- Prolonged mechanical stress inadvertently predisposes a touch-acclimated Arabidopsis plant to infection by the hemibiotrophic fungal pathogen Fusarium oxysporum f. sp. conglutinance (Foc).
- Mechanical stress heightens jasmonic acid and defense gene expression in touch-acclimated plants during Foc infection.
- Susceptibility of touch-acclimated plants to Foc infection was dependent on jasmonic acid biosynthesis mediated by the rate-limiting enzyme JASMONATE RESISTANT 1.
- Prolonged mechanical stress altered the basal levels and MIS-induction of jasmonic acid signalling and defense gene expressions, as well as JA hormone levels in touch-acclimated plants.
- Prolonged MIS primes a yet-to-be identified JA signalling network in touch-acclimated plants that persists without stress to promote susceptibility to the devastating crop pathogen Foc.

## Introduction

In plants, prolonged events of mechanical-induced stress (MIS) cause thigmomorphogenesis, an acclimation phenomenon characterized by reduced petiole length, leaf area, and plant height, along with cell wall thickening and overall enhanced tensile strength (Brenya et al., 2022, Braam, 2005, Börnke and Rocksch, 2018). MIS induces the biosynthesis of chemical compounds that strengthen defense against insect herbivory and necrotrophic pathogen infection (Cipollini, 1997, Zhao et al., 2005, Benikhlef et al., 2013, Markovic et al., 2019, Douma and Anten, 2019, Brenya et al., 2020). For example, Arabidopsis plants exposed to prolonged MIS show transient induction of jasmonic acid (JA) levels in foliar tissues within 10 to 30 min post-stimulation (Brenya et al., 2020, Van Moerkercke et al., 2019). Some pathogens can exploit stress-induced changes in phytohormones, secondary metabolites, and enzyme levels to increase their virulence, bypass host defenses, and successfully invade host tissues (Patkar and Naqvi, 2017, Ma and Ma, 2016, Spanu and Panstruga, 2017, Vargas et al., 2012, Chowdhury et al., 2017). For instance, enhanced JA biosynthesis and signaling can antagonize salicylic acid (SA)-mediated defense, thereby promoting resistance against necrotrophic fungal infection that typically thrives on necrotic plant tissues (Thomma et al., 1998, Glazebrook, 2005, Spanu and Panstruga, 2017, Rabe et al., 2013, Trusov et al., 2009, Thatcher et al., 2009, Rahman et al., 2012). Stress-induced changes that alter JA and/or SA levels can influence plant immunity either positively or negatively, depending on whether the fungal pathogen is biotrophic, necrotrophic, or hemibiotrophic.

The soil-borne fungus *Fusarium oxysporum* is a widespread hemibiotrophic phytopathogen capable of causing significant crop damage and reduced production. More than 106 characterized plant pathogenic forms (*forma specialis*; f. sp.) of *F. oxysporum* have been identified based on their primary host plants (Edel-Hermann and Lecomte, 2018). *Fusarium oxysporum* infiltrates the roots, colonizes the host through the vascular tissue, and initially remains asymptomatic during the biotrophic phase of infection (Lyons et al., 2015). It blocks the host vascular system, causing symptoms such as leaf chlorosis, vein clearing, stunted growth, wilting, poor reproductive development, root rot, and ultimately host death. Managing *F. oxysporum* disease is challenging due to its complex infection strategies, the transition from the biotrophic to necrotrophic phase, persistence in the soil, broad host range, and the exacerbating effects of abiotic environmental stressors (Husaini et al., 2018, Michielse and Rep, 2009, Edel-Hermann and Lecomte, 2018). In Arabidopsis, the three f. sp. of *F. oxysporum* (*F. oxysporum* – f. sp. *conglutinans*; Foc, - f. sp. *raphanin;* FOR and - f. sp. *matthioli,* FOM) have been identified to cause disease via three phases of host interaction; in the prevascular and vascular phases, the pathogen penetrates the roots and enters the xylem vessels, respectively, where it multiplies and develops below ground disease symptoms. In the final expressive phase, disease symptoms become visible in the leaves (Diener, 2012).

Some strains of *F. oxysporum* utilise host hormonal imbalances to infect host cells (Cole et al., 2014, Thatcher et al., 2009, Haidoulis and Nicholson, 2020). In fungal culture filtrates, *F. oxysporum* strains, such as FOM and Foc, can produce jasmonates, including JA, JA-leucine (JA-Leu), and JA-isoleucine (JA-Ile) amino acid conjugates, along with intermediates like 12-oxophytodienoic acid. These fungal-derived jasmonates act as effectors, facilitating root infection through CORONATINE INSENSITIVE1 (COI1)-mediated signaling pathways (Cole et al., 2014, Miersch et al., 1999, Oliw and Hamberg, 2017). Several studies have shown that exogenous application of methyl jasmonate (MeJA) at either the early or late stages of infection compromise and even enhance resistance to *Fusarium* species in various plants (Haidoulis and Nicholson, 2020, Ameye et al., 2015, Makandar et al., 2010, Cole et al., 2014). Emerging evidence indicates that prolonged mechanical-induced stress (MIS) to Arabidopsis primes a touch-acclimated plants to exhibit heightened JA- and defense-related gene expression and immunity to necrotrophic pathogen infection (Brenya et al., 2022, Brenya et al., 2020). Yet it remains unclear if the MIS priming of JA biosynthesis can compromise Foc infection in a touch-acclimated plant.

In this study, we investigated the impact of MIS on Arabidopsis immunity against Foc infection in naïve *vs*. touch-acclimated plant tissues. We examined how single *vs*. prolonged events of MIS affect the balance between JA and SA levels and the impact upon defense gene expression. We hypothesized that touch-acclimated plants would exhibit increased susceptibility to Foc infection. We utilised the JA biosynthesis *jasmonate resistance 1* (*jar1-11*) mutant as well as exogenous phytohormone application to contrast how MIS impacts Foc infection in both naïve and touch-acclimated tissues. These findings provide new mechanistic insights into how MIS affects Arabidopsis immunity against a hemibiotrophic fungal pathogen that causes vascular wilt in closely related cruciferous crops like cabbage, broccoli, and canola.

## Materials and Methods

### Germplasm and plant growth conditions

The following homozygote mutant germplasms were obtained from the Arabidopsis Biological Resource Centre was in the Col-0 ecotype wildtype background; JA biosynthetic mutant *jar1-11* (AT2G46370, CS67935, Salk_034543) showed severe reduction in jasmonyl-L-isoleucine (JA-Ile) accumulation following wounding and decreased sensitivity to MeJA-induced root growth inhibition (Suza and Staswick, 2008, Staswick et al., 1992), SA signalling mutant *npr1-5* (AT1G64280, CS3724); (Greenberg et al., 1994), *grx480* (AT1G28480, SALK_031817C). Homozygous mutant plants were screened by phenotype, antibiotic selection, and/or PCR to confirm T-DNA insertions or point mutations. Primers are displayed in Supplementary Table 1 and 2. Seeds were stratified, and plants were sown in seed-raising soil mixture (DEBCO Pty, Australia) as described (Brenya et al., 2020).

### Mechanically induced stress treatments

Mechanically induced stress (MIS) was performed as described (Brenya et al., 2020). Naive juvenile seedlings (10 days after germination; DAG) and adult rosettes (17 DAG) were gently stimulated in clockwise and counterclockwise motions for 8–10 seconds using a soft brush, constituting a single event of touch stimulation. To generate a touch-acclimated plant, 10-day-old seedlings were touch-stimulated twice daily (8-hour intervals) for 7 days of MIS unless otherwise stated. After 7 days of MIS at 17 DAG, thigmomorphogenesis was quantified in touch-acclimated and naive plants by phenotyping traits such as average rosette area, petiole length, and inflorescence stem height using ImageJ (https://imagej.nih.gov/ij/download.html). Unless otherwise stated, tissue samples were collected 30 minutes following a single MIS event to naïve and/or touch-acclimated plants for further hormone and/or gene expression analysis.

### Foc inoculation and disease assessment

*Fusarium oxysporum* f. sp. *conglutinans* (Foc; BRIP 5176) was obtained from the Victorian Department of Primary Industries in Australia (http://www.dpi.vic.gov.au). Culturing Foc was performed as previously described (Trusov et al., 2009). In brief, underground roots of both touched-acclimated (7 days of MIS) and naïve plants (17 DAG) were inoculated with 500 µL of Foc spore suspension (4 × 10⁶ spores/mL) or water (mock control). Plants were maintained at 28°C under a 12/12 h light/dark photoperiod and covered with a transparent plastic dome. Disease symptoms, characterized by yellow-veined leaves, were scored 8 days after Foc inoculation. Dead plants were identified as those lacking any green leaf material. Disease progression was assessed by counting leaves displaying distinct vein chlorosis or fully yellow leaves. The percentage of diseased plants was calculated as the number of diseased leaves per plant divided by the total number of leaves per plant.

### Foc DNA quantification

To analyse Foc proliferation by DNA copy number in Arabidopsis, DNA was extracted (GeneJET Plant Genomic DNA Purification Kit, Thermo Fisher Scientific, USA) from leaves 6-8 days post-inoculation. DNA was extracted from Foc mycelia. qPCR was used to amplify the MATING-TYPE (MAT1) fungal marker gene (Lievens et al., 2007) in a 20 μl reaction mixture containing either pure *F. oxysporum* genomic DNA (0.1-20 ng) or 100 ng of genomic DNA from infected plants using the Light Cycler 480 SYBR Green 1 Master Mix (Roche, Switzerland) as manufacturer’s instructions. A standard curve was plotted from 10-fold serial dilutions of pure fungal genomic DNA versus threshold cycle (Ct) values from which DNA copy number was determined as described (Lau et al., 2014, Scarlett et al., 2013).

### RNA extraction, cDNA synthesis, and quantitative real-time PCR (qRT-PCR)

Total RNA was isolated from leaf tissue using Meridian Bioscience RNA Extraction Kit according to the manufacturer’s instructions, followed by DNase treatment. to remove residual genomic DNA. RNA concentration and quality were assessed using a DeNovix DS-11 FX spectrophotometer (DeNovix Inc., Wilmington, DE, USA). First-strand cDNA was synthesised using the SensiFAST™ cDNA Synthesis Kit (Meridian Bioscience, Bioline) according to the manufacturer’s instructions. Quantitative real-time PCR (qRT-PCR) was performed on a LightCycler® 480 Instrument (Roche Diagnostics, Switzerland) using FastStart DNA Master SYBR Green I chemistry according to the manufacturer’s instructions. Primer design, validation, and amplification efficiency were optimised as previously described (Brenya et al., 2020). Amplification was conducted under the following cycling conditions: an initial denaturation step at 98°C for 30 s, followed by 30 cycles of 98°C for 10 s, 60°C for 30 s, and 72°C for 30 s, with a final extension at 72°C for 5 min. Relative transcript abundance was calculated using the Pfaffl method (Pfaffl, 2001), which incorporates amplification efficiencies for both target and reference genes. Melting curve analysis was conducted to confirm amplification specificity. Gene expression data were normalised against *ACTIN2* (AT3G18780) as the primary reference gene, and expression stability was validated using *PROTEIN PHOSPHATASE 2A* (*PP2A*; AT1G13320) (Czechowski et al., 2005). Primer sequences used in this study are provided in Table S2.

### Histochemical foliar assays

To examine necrosis, rosette leaves were stained with lactophenol-trypan blue solution (20 mL phenol, 20 mL lactic acid, 40 mL glycerol, 20 mL water and 0.05% trypan blue mix and 200 mL 96% ethanol) for 2 min and de-stained in chloral hydrate (2.5 g in 1 mL of 30% glycerol) overnight. Leaves were viewed under a Carl Zeiss Axio Scope A1 microscope (Carl Zeiss, USA) equipped with interference or phase-contrast optics and a digital camera. Callose deposition was determined by using 95–100% ethanol to remove chlorophyll. After 24 h, leaves were washed in 0.07M phosphate buffer (pH 9.0) and then incubated in fresh buffer containing 0.05% aniline blue (Sigma) for 2 h in the dark. Samples were visualized and photographed with Carl Zeiss Axio Scope A1 microscope equipped with fluorescent UV light (excitation 340 to 380 nm, emission 425 nm).

### Phytohormone treatments

For exogenous treatment with hormones, plants were gently sprayed with a fine mist of salicylic acid (SA, 100 μM), methyl jasmonate (MeJA, 100 μM), or the combination of SA+MeJA (100 μM each) every 2 days as reported (Figure S1A). For control, plants were sprayed with water. Leaves were sampled 30 min after a single touch stimulation event at the end of the spray period (Figure 1A). Rosette area was scored 7 days after the MeJA spray period. Phytohormones were obtained from Sigma (http://www.Sigmaaldrich.com). For the Foc experiments, exogenous MeJA (50 μM) was applied by spraying plants 24–36 hrs post-Foc inoculation.

**Figure 1.**
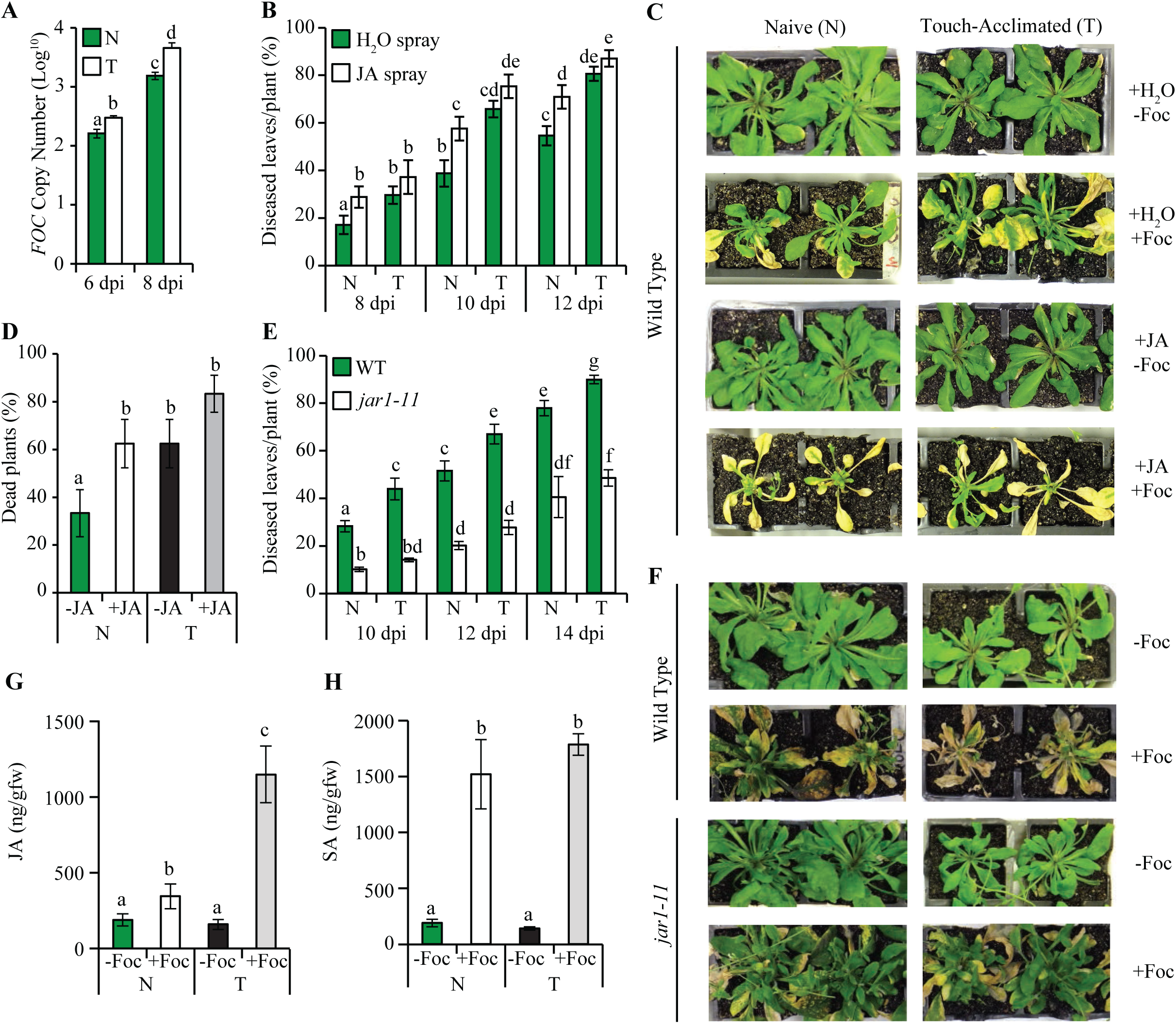
Quantification of Foc infection and phytohormone levels in naïve, touch-acclimated, and MeJA treated WT and *jar1-11* plant tissues. Juvenile seedlings were subjected to seven days of mechanical induced stress generating a touch-acclimated plant at 17 days after germination (DAG). The roots of unstressed (naïve; N) and touch-acclimated (T) plants at 17 DAG were inoculated with (+) or without (-) *Fusarium oxysporum* f. sp. *conglutinans* (Foc). A) Quantification of Foc DNA copy number in naive and touch-acclimated plant leaves at 6- and 8-days post-inoculation (dpi) (n = 3). B) Percentage of diseased leaves per plant showing chlorosis at 8, 10, and 12 dpi in naive and touch-acclimated plants, with (+) or without (-) exogenous methyl jasmonate (JA) spray treatment (n = 10–15). C, F) Representative images of naïve and touch-acclimated WT (C, F) and *jar1-11* (F) plants exposed to various treatments (±Foc, ±JA) at 12-14 dpi. D) Percentage of naive and touch-acclimated plants appearing dead at 15 dpi (± water, ±JA; n = 5–15). E) Percentage of diseased leaves from naive and touch-acclimated wild type (WT) and *jar1-11* (F) plants at 10, 12, and 14 dpi (n = 12). G–H) Quantification of foliar jasmonic acid (JA) and salicylic acid (SA) levels (ng/gfw) in leaves from naive and touch-acclimated plants with (+) or without (-) exposure to Foc infection (5-days post-inoculation). Letters are statistical differences between treatments using ANOVA with the Bonferroni test (*p* < 0.05).

### Phytohormone quantification

For quantifying JA, JA-ILE and SA, 3-5 replicates of leaf samples (∼200mg) were grounded to fine powder. Hormones were extracted using 1ml of extraction solvent containing Methanol: water:acetic acid, 69:30:1 (v/v) and 100ppb of deuterated phytohormones (Olchemim Ltd, Olomouc, Czech Republic) as internal standards. The blank control contained extraction solvent with the internal standard only. Samples for vortexed briefly, placed in sonicator for 30 minutes at 4-8 °C and centrifuged at 15,000 rpm for 10 minutes at 4 °C. The supernatant was transferred into a new eppendorf tube, evaporated to dryness and re-suspended in 200 μL 70% methanol (v/v) by sonicating for 10 minutes at 4 -8 °C. Finally, samples were centrifuged at 15,000 rpm for 10 minutes and 50 μL of the supernatant solution was transferred into chromatography sample plates for LC-MS. The chromatographic separation was achieved using Waters ACQUITY UPLC I-Class system (Waters Corporation, Milford, MA, USA) equipped with ACQUITY UPLC BEH C18 Column, 130Å, 1.7 µm, 2.1 mm X 50 mm column, set at 45 °C. The mobile phase consisting of H_2_O with 0.1% formic acid (mobile phase A) and Acetonitrile with 0.1% formic acid (mobile phase B) were operated under a gradient elution program and flow rate of 0.6ml/minute. Detection of ABA and JA was carried out in negative electron spray ionization mode and multiple reaction monitoring in SCIEX Triple QuadTM 7500 LC-MS/MS system at the Western Sydney University Mass Spectrometry Facility, Campbelltown, Australia. A calibration curve was created for quantification containing pure unlabelled compound (SA and JA) from 0.061 ppb to 250 ppb and 100 ppb of internal standard. The pure JA standard was obtained from Sigma-Aldrich. Internal standards included deuterated [²H₄]-SA (C₇H₂D₄O₃) from Olchemim Ltd. (Olomouc, Czech Republic) and jasmonic-d₅ acid (C₁₂H₁₃D₅O₃) from CND Isotopes (Canada).

### Statistical analysis

Data was statistically analysed using R, JMP Pro software (Version 19; SAS Institute Inc., Cary, NC, USA) or SigmaPlot 14.0 (Version 19). Data were first assessed for normality and homogeneity using of variance (biological replicates) prior to statistical analysis. Differences among treatment groups were evaluated using one-way analysis of variance (ANOVA). When a significant treatment effect was detected (P < 0.05), means were separated using Tukey’s Honestly Significant Difference (HSD) or Bonferroni post hoc test at the 5% significance level. For comparisons involving only two groups, Student’s t-test was used. Data are presented as mean ± standard error (SE), and differences were considered statistically significant at *p* < 0.05.

## Results

### Prolonged MIS hyper-induces JA in Arabidopsis to promote susceptibility to Foc

Ten-days-old juvenile Arabidopsis plants were exposed to 10 seconds of mechanical induced stress (MIS) twice daily for 7-days generated a touch-acclimated plant 17 days after germination (DAG) (Figure S1A). The touch-acclimated adult plants exhibited a classic thigmomorphogenesis phenotype characterised by a reduction rosette area (50%) and shorter floral stem height that persisted for 28 (66%) and 41 (23%) DAG (Figure S1B-C). To investigate the impact of prolonged MIS on plant immunity against the hemibiotrophic pathogen *Fusarium oxysporum* f. sp. *conglutinans* (Foc), the control and touch-acclimated plant (17 DAG) were root inoculated with 500 μl fungal spores (4 x 10^6^ spores/ml) or water. To monitor the systemic progression of Foc from the roots to the shoots, we quantified fungal DNA in the leaves, which allowed us to evaluate how rapidly the infection spread to aerial rosette tissues - the direct targets of our abiotic stress treatment. After 6- and 8-days post-inoculation (dpi), there was a significant increase in copy number of the *MATING-TYPE* (*MAT1*) gene in touch-acclimated compared to naïve control plants, reflecting that MIS enhances fungal development (Figure 1A). Staining of leaves at 7 dpi from touch-acclimated plants with trypan blue or aniline blue showed no significant difference in necrotic lesions or chitin-triggered callose deposition, respectively to the naïve control plants (Figure S1D-E). Disease progression and symptoms, including yellowing of leaf veins (chlorosis) and tissue decay, were assessed as previously described (Trusov et al., 2009). Symptoms became evident around 8 dpi, marking the onset of the necrotrophic phase. Compared to naïve plants, the touch-acclimated plants exhibited significantly more chlorotic and diseased leaves (>25%) between 8 and 12 dpi (Figure 1B-C). By 15 dpi, 60% of MIS plants had died, lacking visible green rosette leaves, compared to 35% of naïve plants, demonstrating that prolonged MIS makes plants highly susceptible to Foc infection (Figure 1D).

Prolonged MIS promotes thigmomorphogenesis via JA biosynthesis and signalling (Chehab et al., 2012). We hypothesised that prolonged MIS would heighten JA biosynthesis/signaling thereby interfering with defense against Foc. Plants sprayed with a fine mist of water every other day for 7 days exhibited no change in rosette size or inflorescence stem height at 17 DAG compared to untreated control plants (Figure S1). Exogenous application of 100 μM methyl jasmonate (JA) reduced rosette area (54%) and inflorescence stem height (77% at 28 DAG, and 34% at 41 DAG) relative to control plants, thereby mimicking touch-induced thigmomorphogenesis (Figure S1A-C). Compared to water (H_2_O) sprayed plants, exogenous spraying with JA in the absence of any MIS exhibited a higher percentage of diseased leaves between 8 to 12 dpi with Foc (Figure 1B-C). No significant differences in disease susceptibility to Foc were observed between JA-treated (-MIS) and touch-acclimated (+MIS) plants (Figure 1B-C). By 15 dpi, 63% of JA-treated and touch-acclimated plants appeared brown and dead (Figure 1D). Therefore, JA treatment mimics prolonged MIS by promoting thigmomorphogenesis and Foc susceptibility in the acclimated plant.

To test whether MIS primes JA responses and susceptibility to Foc, we used the JA biosynthesis mutant *jar1-11,* previously shown to impair thigmomorphogenesis (Chehab et al., 2012). Compared with wild-type (WT) plants, *jar1-11* mutant plants exhibited ∼50% less Foc disease symptoms throughout infection (10-14 dpi; Figure 1E, F). Unlike the WT touch-acclimated plants, which showed significantly higher susceptibility to Foc than WT untouched control, *jar1-11* touch-acclimated plants exhibited disease symptoms similar to *jar1-11* control plants at 10-, 12- and 14 dpi (Figure 1E-F). The MeJA-induced susceptibility to Foc prompted us to examine JA and SA levels in leaves following a single touch stimulation event in naïve (control) and touch-acclimated (+MIS) plants, with (+) or without (-) Foc infection. After 5 dpi, JA levels increased by 0.82-fold in control Foc-infected plants compared to uninfected control plants (Figure 1G). In the touch acclimated-infected plants, JA level was significantly induced up to 6.17-fold compared to uninfected touch-acclimated plants. In contrast, SA levels were induced by Foc infection at similar levels in both control and touch-acclimated leaf tissues (Figure 1H). Overall, prolonged MIS primes the touch-acclimated plant such that during Foc infection JA levels become heightened, enabling Foc host susceptibility and increased disease symptoms.

### Prolonged MIS hyper-induces defense gene expression during Foc infection in a JA-dependent manner

Prolonged MIS was shown to heighten JA biosynthesis and defense gene expression during necrotrophic pathogen infection of a touch-acclimated relative to a naive plant (Brenya et al., 2020, Van Moerkercke et al., 2019). We investigated the interaction between prolonged MIS and Foc infection by quantifying the expression of the JA biosynthetic gene *OXOPHYTODIENOATE-REDUCTASE 3* (*OPR3*), the JA-signaling regulator *MYELOCYTOMATOSIS ONCOGENE HOMOLOG 2* (*MYC2*), and the JA-responsive defense marker genes *VEGETATIVE STORAGE PROTEIN 1* (*VSP1*), and *PLANT DEFENSIN 1.2* (*PDF1.2*) in leaves from naïve and touch-acclimated plants 3 and 5 dpi with or without Foc. The relative expression of all four genes was significantly higher after 3 dpi in touch-acclimated plants (Figure 2A). That is, their foliar expression became hyper-induced in infected touch-acclimated relative to infected naive plants, ranging from 4.0- to 12.1-fold for *OPR3*, 3.5- to 8.2-fold for *MYC2*, 6.3- to 47.2-fold for *VSP1*, and 9.2- to 23.4-fold for *PDF1.2* (Figure 2A). Defense gene expression in infected naïve plants tended to be higher than the uninfected control, however only *PDF1.2* expression was significantly induced by 9.3-fold. At 5 dpi, expression of all four genes increased 5.6- to 108-fold in leaves of infected relative to uninfected naive plants (Figure 2B). The hyper-induction of *OPR3* (6.3- to 7.7-fold), *MYC2* (5.6- to 14.5-fold), *VSP1* (70.6- to 240.7-fold), and *PDF1.2* (108-to 246-fold) remained elevated at 5 dpi in infected touch-acclimated compared to naive plants. Intriguingly, the basal level of defense gene expression in uninfected touch-acclimated plants remained similar to the naïve control, except for *VSP1,* which was constitutively 25.7- to 41.4-fold higher at 3 and 5 days post recovery after ceasing MIS events (Figure 2A-B). Thus, prolonged MIS primes defense gene expression during Foc infection of a touch-acclimated plant.

**Figure 2.**
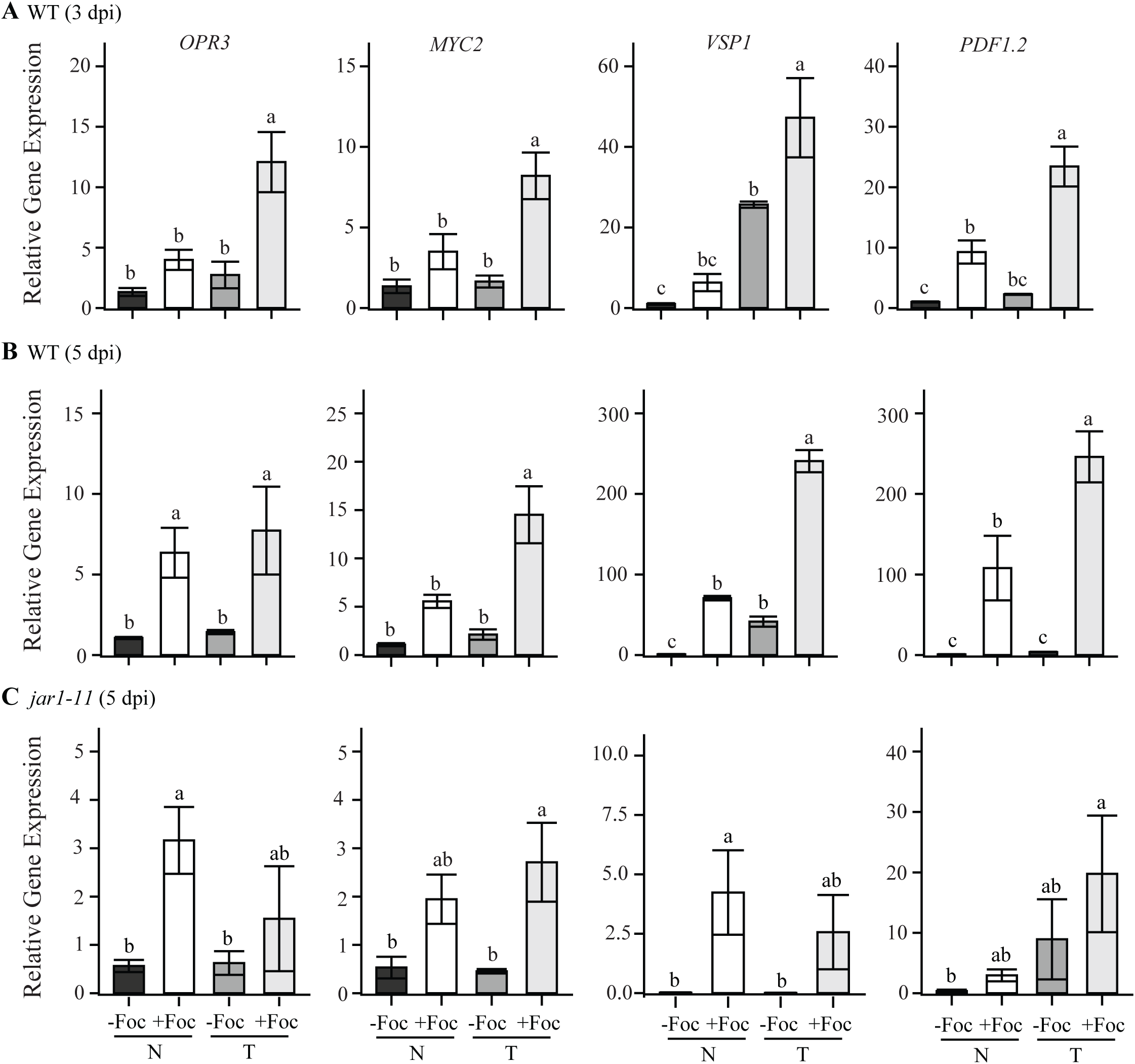
Quantification of JA- and defense-responsive gene expression in Foc-infected foliar tissues from naïve and touch-acclimated plants. Juvenile seedlings were subjected to seven days of mechanical induced stress generating a touch-acclimated plant at 17 days after germination (DAG). The roots of unstressed (naïve; N) and touch-acclimated (T) plants at 17 DAG were inoculated with (+) or without (-) *Fusarium oxysporum* f. sp. *conglutinans* (Foc). A-C) Relative transcript levels of JA- and defense-responsive genes in (A) wild type (WT) plants at 3 dpi, (B) WT plants at 5 dpi, (C) *jar1-11* mutant at 5 dpi (expression is relative to the naïve WT control plant). The expression of genes were normalized against the *ACTIN 2* housekeeping gene. Error bars show standard error (n=3). Data is a representative of multiple independent experiments. Letters are statistical differences between treatments using ANOVA with the Bonferroni test (*p* < 0.05). *PDF1.2; PLANT DEFENSIN 1.2*, *VSP1*; *VEGETATIVE STORAGE PROTEIN 1*, *MYC2; MYELOCYTOMATOSIS ONCOGENE HOMOLOG 2*, *OPR3; OXOPHYTODIENOATE-REDUCTASE 3*.

To determine if the priming of JA- and defense-related gene expression in infected touch-acclimated plant tissues was dependent on JA, we quantified transcript levels in the *jar1-11* mutant at 5 dpi (Figure 2C). Unlike WT, the Foc-infected tissues from naïve and touch-acclimated plants *jar1-11* plants displayed no significant differences in gene expression. The *jar1-11* Foc-infected naïve plant tissues exhibited a modest yet significant upregulation of *OPR3* (3.2-fold) and *VSP1* (4.3-fold) relative to the uninfected naïve control. Compared to WT (Figure 2B), the overall expression of *OPR3, VSP1, MYC2* and *PDF1.2* were vastly diminished in *jar1-11* (Figure 2C). The basal expression of all genes (including *VSP1*) in *jar1-11* touch-acclimated and naïve leaf tissues remained unchanged. Therefore, JAR1 is required to enhance Foc induction and hyper-induction of JA- and/or defense-responsive gene expression in naïve and touch-acclimated plant tissues respectively.

### Mechanical stress regulation of defense gene expression in naïve and touch-acclimated tissues

We next investigated whether prolonged MIS can hyper-induce the expression of defense genes in touch-acclimated plant tissues following a single MIS event in the absence of Foc infection. The regulation of stress responsive (*RD29A; RESPONSIVE TO DESICCATION 29A, RD20*; *RESPONSIVE TO DESICCATION 20*), JA biosynthetic (*AOS; ALLENE OXIDE SYNTHASE, OPR3; 12-OXO-PHYTODIENOIC ACID REDUCTASE*), JA signalling (*MYC2; MYELOCYTOMATOSIS ONCOGENE HOMOLOG 2; LOX1; LIPOXYGENASE 1*), JA/defense responsive (*PDF1.2; PLANT DEFENSIN 1.2, VSP1; VEGETATIVE STORAGE PROTEIN 1,*) and SA/defense responsive (*NPR1; NONEXPRESSER OF PATHOGENESIS-RELATED GENES 1, GRX480*; *GLUTAREDOXIN 480*) genes were quantified in leaves from naïve and touch-acclimated plants (10 DAG and 17 DAG), 30 min before and after a single event of touch stimulation.

The expression of general stress responsive (*RD29A* and *RD20*) and JA biosynthetic structural genes (*OPR3* and *AOS*) were all similarly induced 2.2 to 22.3-fold by MIS in leaves of naive juvenile plants and naïve and touch-acclimated adult plants (Figure 3A-B). The expression of JA signalling genes (*MYC2* and *LOX1*) was significantly induced (2.5 to 3.5-fold) by MIS in adult naïve but not in touch-acclimated plants. *MYC2* was also significantly induced in naïve juvenile plant by 11.2-fold following MIS (Figure 3C). The expressions of JA and defense responsive genes, *VSP1* and *PDF1.2*, exhibited opposite trends compared to JA biosynthetic genes. Expression of these genes declined - 1.5 to -3.5-fold following MIS in adult naïve and touch-acclimated tissues, yet only *PDF1.2* was significantly suppressed. In naïve juvenile plants, MIS suppressed *VSP1* expression, whereas *PDF1.2* remained unchanged.

**Figure 3.**
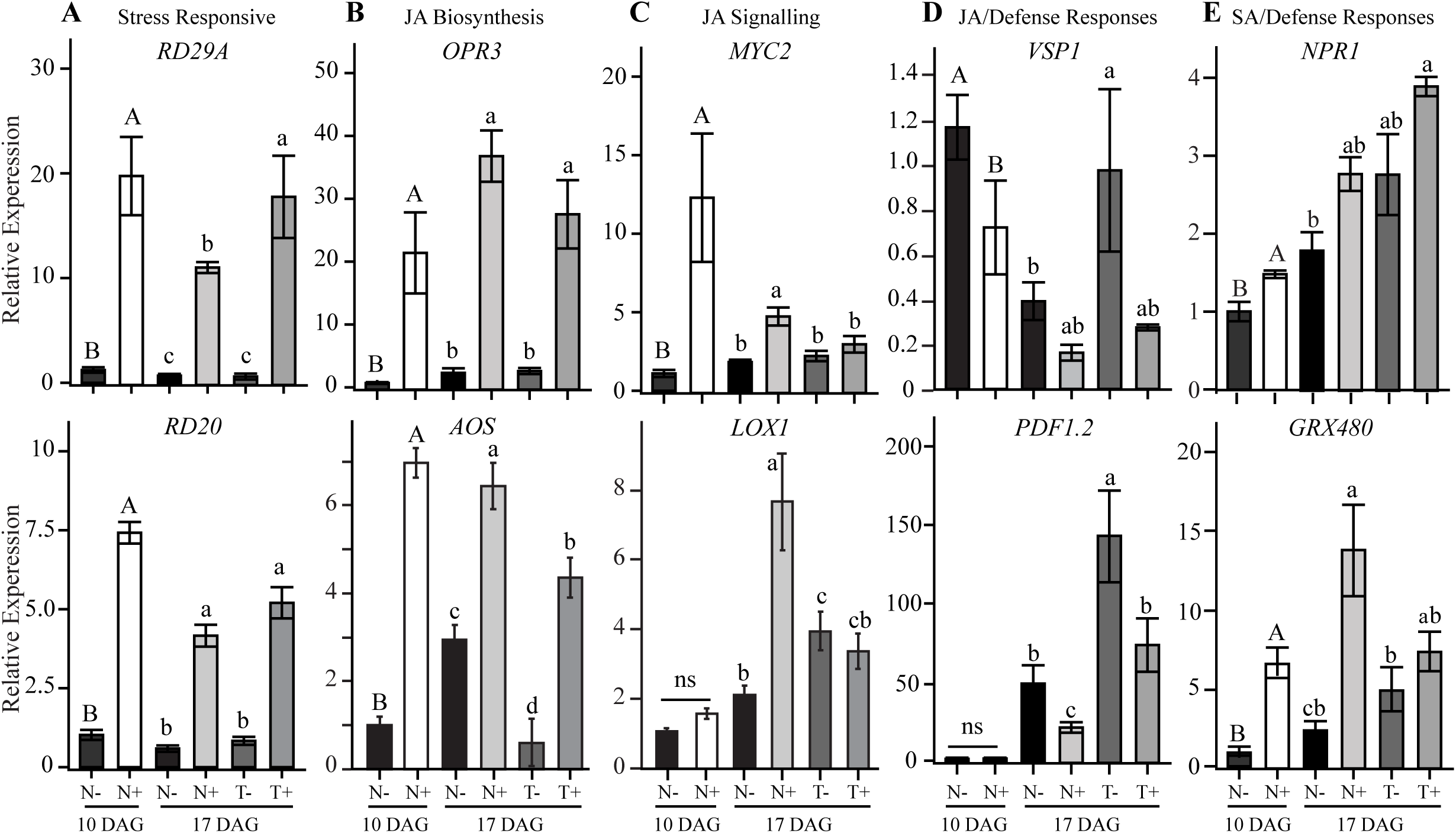
Quantification of stress, JA biosynthesis, signaling, and defense-related gene expression in naïve and touch-acclimated foliar tissues before and after MIS. Juvenile (10 days after germination; DAG) seedlings were subjected to seven days of mechanical induced stress (MIS) generating a touch-acclimated plant at 17 DAG. Gene expressions were quantified in juvenile and adult naive (N) and adult touch-acclimated (T) plant tissues before (-) and 30 min after a single event of MIS (+). The expression levels of genes were normalized to the *ACTIN 2* housekeeping gene. Error bars show the standard error of the mean (n = 4-7). Data is representative of multiple independent experiments. Capital and lower-case letters respectively define statistical differences between treatments using ANOVA (10 DAG) and Bonferroni test (17 DAG). *RD29A; RESPONSIVE TO DESICCATION 29A, RD20*; *RESPONSIVE TO DESICCATION 20*, *AOS; ALLENE OXIDE SYNTHASE, OPR3; 12-OXO-PHYTODIENOIC ACID REDUCTASE*, *MYC2; MYELOCYTOMATOSIS ONCOGENE HOMOLOG 2; LOX1; LIPOXYGENASE 1*, *PDF1.2; PLANT DEFENSIN 1.2, VSP1; VEGETATIVE STORAGE PROTEIN 1, NPR1; NONEXPRESSER OF PATHOGENESIS-RELATED GENES 1, GRX480*; *GLUTAREDOXIN 480*).

The SA signalling gene *NPR1* was induced by MIS only in naïve tissues, whereas *GRX480* was significantly up-regulated (∼5.6- to -6.1-fold) following MIS in both naïve and adult plants but not in touch-acclimated plants (Figure 3E). The GRX480 expression pattern was opposite of *PDF1.2* expression pattern which confirms with previous reports that *GRX480* is a negative regulator of *PDF1.2* (Ndamukong et al., 2007). Finally, the basal expression levels of *LOX1, VSP1* and *PDF1.2* were significantly higher in touch-acclimated relative to naïve adult plants prior to MIS (Figure 3C-D).

In summary, general stress responsive and JA structural biosynthetic genes were positively regulated by MIS, whereas JA/defense-related genes were repressed by MIS in adult naïve and touch-acclimated plant tissues. JA signalling genes were touch-induced in adult naïve tissues but lost their induction response in touch-acclimated tissues. Some genes involved in JA signaling and defense showed higher basal expression levels in touch-acclimated tissues. Therefore, unlike the hyper-induction of transcriptional responses observed in touch-acclimated plants during Foc infection (Figure 2), a single event of MIS in uninfected touch-acclimated plants did not heighten the transcriptional response of the suite of genes tested.

### MIS induces JA but not SA in naïve and touch-acclimated plant tissues

We investigated whether prolonged MIS could prime the hyper-induction of JA and SA levels in touch-acclimated plant tissue following a single MIS event in the absence of Foc infection. Across multiple experiments, both naïve juvenile and adult plant tissues showed a consistent increase in JA and JA-Ile levels 2.3- to 60.3-fold relative to control plants following a single MIS event of 10 seconds (Figure 4A-B). Whereas in touch-acclimated plants, while there was a trend showing JA-Ile and JA increased slightly following MIS, yet JA-Ile was only significantly induced by 7.6-fold during one experiment. The levels of SA were inconsistent across multiple experiments; in one experiment SA was induced by an event of MIS regardless of plant age and touch-acclimation, whereas in a second experiment there were no significant change in SA following an event of MIS. We have observed this consistent trend of JA induction accompanied by either the induction or lack of change in SA levels across multiple time points during prolonged MIS. Overall, naïve juvenile and adult foliar tissues subjected to MIS exhibit a consistent induction of JA and JA-Ile, that becomes saturated in touch-acclimated plants.

**Figure 4.**
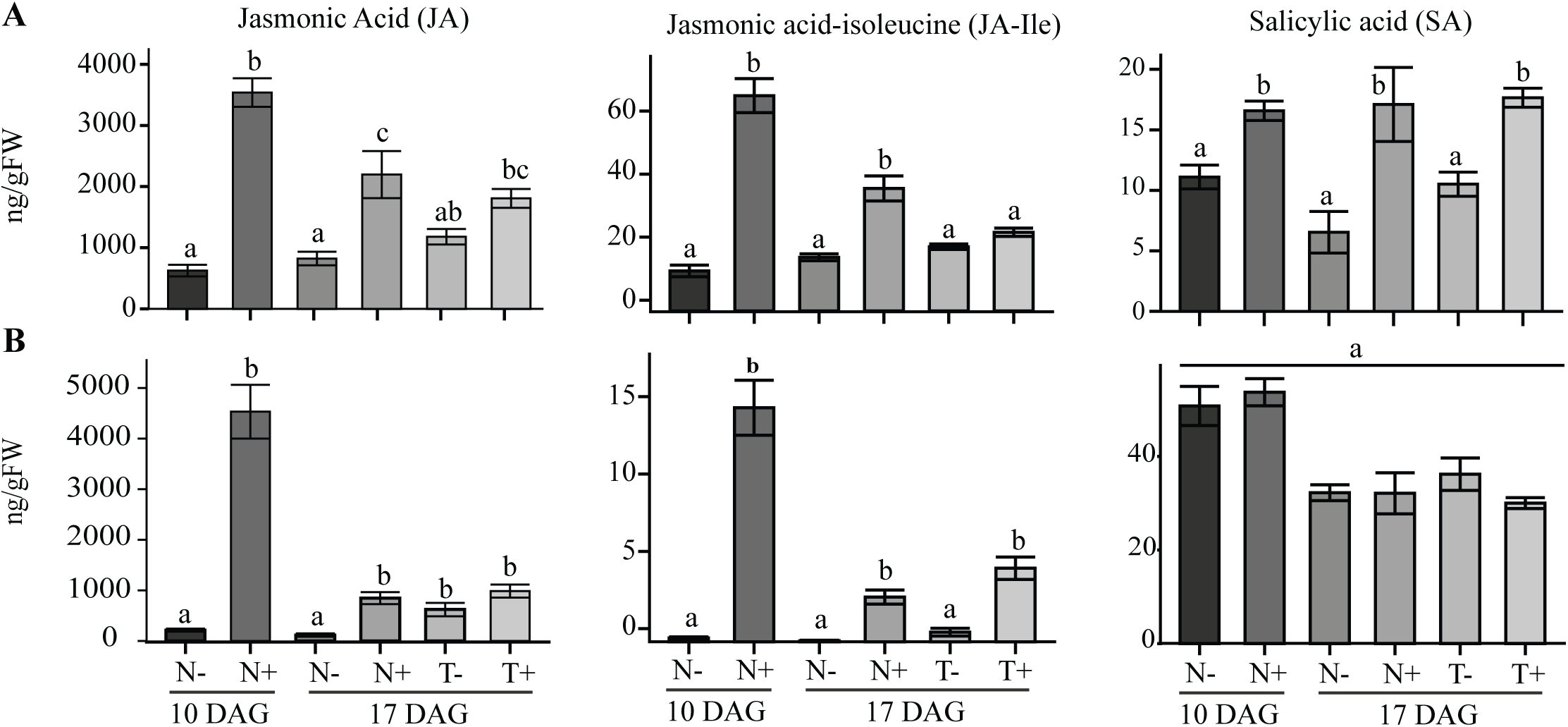
Quantification of phytohormones in naïve and touch-acclimated foliar tissues before and after MIS. Juvenile (10 days after germination; DAG) seedlings were subjected to seven days of mechanical induced stress (MIS) generating a touch-acclimated plant at 17 DAG. Jasmonic acid (JA), jasmonic acid-isoleucine (JA-Ile) and salicylic acid (SA) were quantified in juvenile and adult naive (N) and adult touch-acclimated (T) plant tissues before (-) and 30 min after a single event of MIS (+). A-B) Two independent experiments are shown to highlight consistent and inconsistent changes in JA and SA following MIS. These data sets are representative of multiple independent experiments. Error bars show the standard of the mean (n = 5-8), and letters denote statistical differences between treatments using ANOVA with the Bonferroni test (*p* < 0.05).

## DISCUSSION

Mechanical stimulation enhances JA biosynthesis/signalling and associated gene networks triggering thigmomorphogenesis, a phenotypic phenomenon and the process by which plants acclimate to the cellular vibrations (Van Moerkercke et al., 2019, Chehab et al., 2012, Xu et al., 2018, Lee et al., 2005, Brenya et al., 2022). Consistent with JA’s role in promoting thigmomorphogenesis, exogenous application of MeJA induced a touch-acclimated plant phenotype (Figure S1). Previous work showed that prolonged MIS enhanced the JA-dependent hyper-induction of defense gene expression that correlated with resistance against necrotrophic pathogens such *Botrytis cinerea* and *Alternaria brassicicola* (Brenya et al., 2020). Yet the effects of prolonged MIS on Arabidopsis immunity to hemibiotrophic fungal pathogens like *Fusarium oxysporum* f. sp*. conglutinans* (Foc) that deploy JA to promote infection remained unknown. Here, we demonstrate that prolonged MIS primes a touch-acclimated Arabidopsis plant to become more susceptibile to Foc infection, evident by the hyper-induction of JA levels and defense-related gene expression during Foc infection. MIS transiently increases JA and jasmonoyl-isoleucine (JA-Ile) in leaves of naïve juvenile and adult plants, and to a lesser extent in touch-acclimated plants. Collectively, these findings reveal that prolonged MIS can cross-prime Foc susceptibility and infection in touch-acclimated plants in a JA-dependent manner.

### Prolonged MIS predisposes Arabidopsis plants to Foc infection

Cell wall reinforcements such as lignin and callose depositions observed in touch-acclimated leaves have been reported to deter necrotrophic pathogens from infection (Chehab et al., 2012, Brenya et al., 2020). Since Foc primarily infects plants through the roots and aggressively colonizes the vascular system, it potentially evades foliar structural defenses induced by prolonged MIS. However, foliar-induced jasmonates act as effectors to promote Foc colonization and infection in plants (Cole et al., 2014, Miersch et al., 1999). JA can be conjugated to isoleucine by JAR1, producing the biologically active form JA-Ile during plant stress events (Miersch et al., 2008). JA-Ile binds to the F-box protein CORONATINE INSENSITIVE 1 (COI1), facilitating its interaction with JASMONATE ZIM-domain (JAZ) transcriptional repressor proteins. This interaction promotes JAZ degradation via the 26S proteasome, thereby activating JA signaling pathways (Zhang et al., 2018). Loss-of-function in *JAZ7* increases JA levels and enhances susceptibility to *F. oxysporum*, whereas mutations in MYC2, a positive regulator of JA signaling, confer resistance to *F. oxysporum* (Anderson et al., 2004, Kidd et al., 2009, Thatcher et al., 2012, Nalam et al., 2015, Thatcher et al., 2016). We hypothesised that prolonged MIS would induce JA and increase susceptibility to Foc infection. Consistent with previous reports, exogenous MeJA application promoted thigmomorphogenesis and enhanced Foc infection (Nalam et al., 2015, Trusov et al., 2009, Makandar et al., 2010, Thatcher et al., 2009). Touch-acclimated plant tissues infected with Foc displayed increased disease symptoms such as leaf decay and vein bleaching, as well as higher foliar Foc DNA copy number (Figure 1). Like *F. oxysporum* f. sp. *mathioli* (Fom), Foc can produce isoleucine- and leucine-conjugated jasmonates (JA-Ile/Leu) that enhanced host colonization (Kidd et al., 2011, Thatcher et al., 2016, Thatcher et al., 2009, Cole et al., 2014). MeJA can be metabolized into JA-Ile, which enhances susceptibility to Fom (Cole et al., 2014, Kazan and Manners, 2008, Tamogami et al., 2012, Nalam et al., 2015). Single events of MIS induced both JA and JA-Ile in naïve and touch-acclimated tissues, and the MIS-induction of JA was exacerbated in touch-acclimated plant tissues infected with Foc (Figures 1 and 4).

Previous studies have shown that mechanically shaking for one minute daily over 21 days can also increase susceptibility to *F. oxysporum* f. sp. *lycopersici* (Fol) in tomato (*Lycopersicon esculentum*), flax (*Linum usitatissimum*), and pea (*Pisum sativum*) plants (Shawish and Baker, 1982). Similarly, wind-induced mechanical stress at 7.3 m/s triggered severe outbreaks of leaf and panicle blast diseases caused by hemibiotrophic pathogens *Pyricularia grisea* and *Magnaporthe grisea* in rice (Taguchi et al., 2014). However, some studies report the protective effects of mechanical stress. For example, koa trees subjected to three weeks of bending showed improved survival rates against *F. oxysporum* f. sp. *Koae* (Fok) (Ishihara et al., 2021). Similarly, air-blasting tomato plants at 4 m/s for 30 minutes reduced root infection by *F. oxysporum* f. sp. *radicis-lycopersici* (Forl) (Elsharkawy et al., 2015). These contrasting outcomes may arise from differences in 1) type, intensity, and duration of MIS, 2) developmental stages and growth conditions, 3) variations in plant species, and 4) *F. oxysporum* formae speciales, which collectively can all impact pathogen infection. Different F. oxysporum formae speciales possess distinct repertoires of virulence effectors and pathogenicity determinants that interact differently with different host resistance and susceptibility mechanisms (van Dam et al., 2016, Jangir et al., 2021). Foc may promote infection by manipulating host JA biosynthesis and/or signalling pathways plants treated with MeJA as well as exposed to prolonged MIS (Thatcher et al., 2009, Manstretta and Rossi, 2016, Edel-Hermann and Lecomte, 2018, Cole et al., 2014). Therefore, prolonged MIS may generate touch-acclimated plants while increasing their susceptibility to FOC infection.

### Foc infection of touch-acclimated plant tissues depends upon JA biosynthesis

Loss-of-function mutations in key JA pathway components including *AOS*, *OPR3*, and the transcription factors, MYC2/MYC3/MYC4, have been reported to impair thigmomorphogenesis (Chehab et al., 2012, Van Moerkercke et al., 2019). Here we showed that the loss-of-function mutation in *JAR1*, a key gene in JA biosynthesis, reduced overall Foc infection in *jar1-11* compared to WT (Figure 1). Additionally, there was no difference in the percentage of diseased leaves per plant between the naïve and touch-acclimated *jar1-11* mutant plants (Figure 2). This implies that JA biosynthesis is a critical mediator of MIS-induced disease susceptibility to Foc. Although JAR1 is required for JA biosynthesis to promote thigmomorphogenesis and Foc infection, stress-responsive and JA biosynthetic genes continue to be induced in *jar1-11* touch-acclimated tissues, albeit to a lesser extent following a single event of MIS. At 3 and 5 dpi, the expression of JA-responsive genes *PDF1.2*, *VSP1*, *MYC2*, and *OPR3* were hyper-induced by a single event of MIS in infected touch-acclimated relative to infected naïve plants. However, the loss-of-function in JAR1 markedly reduced the expression of these JA- and defense-responsive genes during Foc infection of naïve *jar1-11* plants, and their hyper-induction was no longer apparent in touch-acclimated infected *jar1-11* plants. Together, these findings suggest that Foc exploits the JAR1-dependent JA signaling to suppress host defenses and promote infection in touch-acclimated plants.

### MIS induces JA in naïve juvenile and adult touch-acclimated plant tissues

It is often proposed that JA induces resistance against necrotrophic pathogens while SA induces resistance against biotrophic pathogens. Although SA–JA crosstalk is thought to involve an adaptive reciprocal antagonism, this remains speculative (Thaler et al., 2012). The hemibiotrophic pathogen Foc is known to manipulate hormonal imbalances by increasing JA to suppress host defenses (Thatcher et al., 2009, Ma and Ma, 2016). Indeed, there was an obvious hyper-induction of JA in touch-acclimated leaves during Foc infection that was accompanied by increased fungal DNA and disease progression (Figure 1). However, SA levels were comparably elevated in Foc-infected naive and touch-acclimated plants relative to the uninfected naïve plants (Figure 1). Previous studies have shown that both SA and JA accumulate in response to *F. oxysporum* and play a role in effector-triggered immunity (Liu et al., 2016, Makandar et al., 2010). Plants engage the SA pathway to antagonize JA-dependent pathogen strategies. Mutants impaired in SA signaling such as *salicylic acid induction-deficient 2* (*sid2*), *enhanced disease susceptibility 5* (*eds5*), and overexpression of *SALICYLATE HYDROXYLASE* (NahG) which prevents SA accumulation, show heightened susceptibility to Foc infection (Trusov et al., 2009, Diener and Ausubel, 2005, Makandar et al., 2010). Yet, a single event of MIS to a naïve or touch-acclimated plant does not consistently induce SA in naïve or touch-acclimated plants. The antagonism between JA and SA pathways has been questionable following MIS (Makandar et al., 2010). In agreement, our results showed that MIS consistently elevates JA but not necessarily SA levels and this likely heightens Foc susceptibility in Arabidopsis.

### Prolonged MIS cross-primes a stress memory potentially involving a JA signalling network

Stress cross-priming occurs where exposure to a primary stress of one type prepares a plant to mount a stronger or weaker defense against a completely different secondary stress (Brenya et al.). JA plays a critical role in mediating MIS-induced stress tolerance and the touch-acclimated plant phenotype (Van Moerkercke et al., 2019, Chehab et al., 2012, Xu et al., 2018, Lee et al., 2005, Brenya et al., 2020). Here we showed that prolonged MIS primed the touch-acclimated plant to heighten JA levels and JA/defense gene expression during Foc exposure, which elevated infection. The induction of JA and JA-Ile levels seen in naïve plants were sustained to some degree in touch-acclimated plants, as was the expression of genes involved in general stress responsiveness (*RD29A*, *RD20*) and JA biosynthesis (*OPR3*, *AOS*). Yet a stress memory becomes evident when the JA master signalling regulator *MYC2* and suppressor *GRX480* genes became unresponsive to a single event of MIS in touch-acclimated leaves. *GRX480* plays a role in modulating cross-priming JA signalling rather than biosynthesis networks (Ndamukong et al., 2007). Prior reports show that exogenous MeJA application fails to induce *GRX480* expression in the JA-insensitive mutant *jin1-1/myc2* (Köster et al., 2012) and *GRX480* expression is MYC2-dependent (Park et al., 2013, Zander et al., 2020). This could explain why *GRX480* and *MYC2* expression, as well as JA and JA-Ile hormone levels, appear saturated following a single event of MIS in touch-acclimated plants.

The JA-responsive marker gene *PDF1.2* is commonly used to interrogate stress-induced plant immunity and defense mechanisms against pathogens (Memelink, 2009). Our data showed that *PDF1.2* expression in naïve and touch-acclimated tissues depends on both plant age and the duration of mechanical stress. In naïve adult but not juvenile plants, a single MIS event significantly upregulated *GRX480* while repressing *PDF1.2* expression, consistent with previous findings showing that *GRX480* overexpression antagonizes *PDF1.2* (Ndamukong et al., 2007). The antagonism of *PDF1.2* expression by *GRX480* induction was age-dependent, since *GRX480* expression was strongly induced in naïve juvenile tissues, yet *PDF1.2* remained unchanged. This aligns with previous studies reporting age-related modulation of stress-responsive genes (Giuntoli et al., 2017, Li et al., 2023).

Prolonged MIS hyper-induces JA levels and defense gene expression in touch-acclimated plants exposed to Foc. We propose that prolonged MIS primes a yet-to-be identified JA signalling network in touch-acclimated plants that persists without further stress to promote long-lasting immunity to necrotrophs and susceptibility to hemibiotrophs. While MIS can benefit agriculture by promoting floral pollination, increasing crop productivity, and priming resistance to necrotrophs, it can inadvertently predispose plants to early infection by a hemibiotrophic fungal pathogen.

## Acknowledgements

We acknowledge Meena Mikhael and Emma Kuebler from the Western Sydney University Mass Spectrometry Facility for their technical support. AB was awarded a Bangabandhu Science and Technology Fellowship trust scholarship.

## Authors Contributions

EB, CIC, MP, AB conceived ideas and designed experiments. EB, MP, AB, LP performed experiments, data collection, prepared figures and contributed to drafting the paper. CIC supervised EB, AB, and MP. DT and ZC co-supervised EB. SJ co-supervised AB and MP. All authors have read and edited the manuscript.

## SUPPLEMENTARY FIGURE LEGENDS

**Figure S1.**
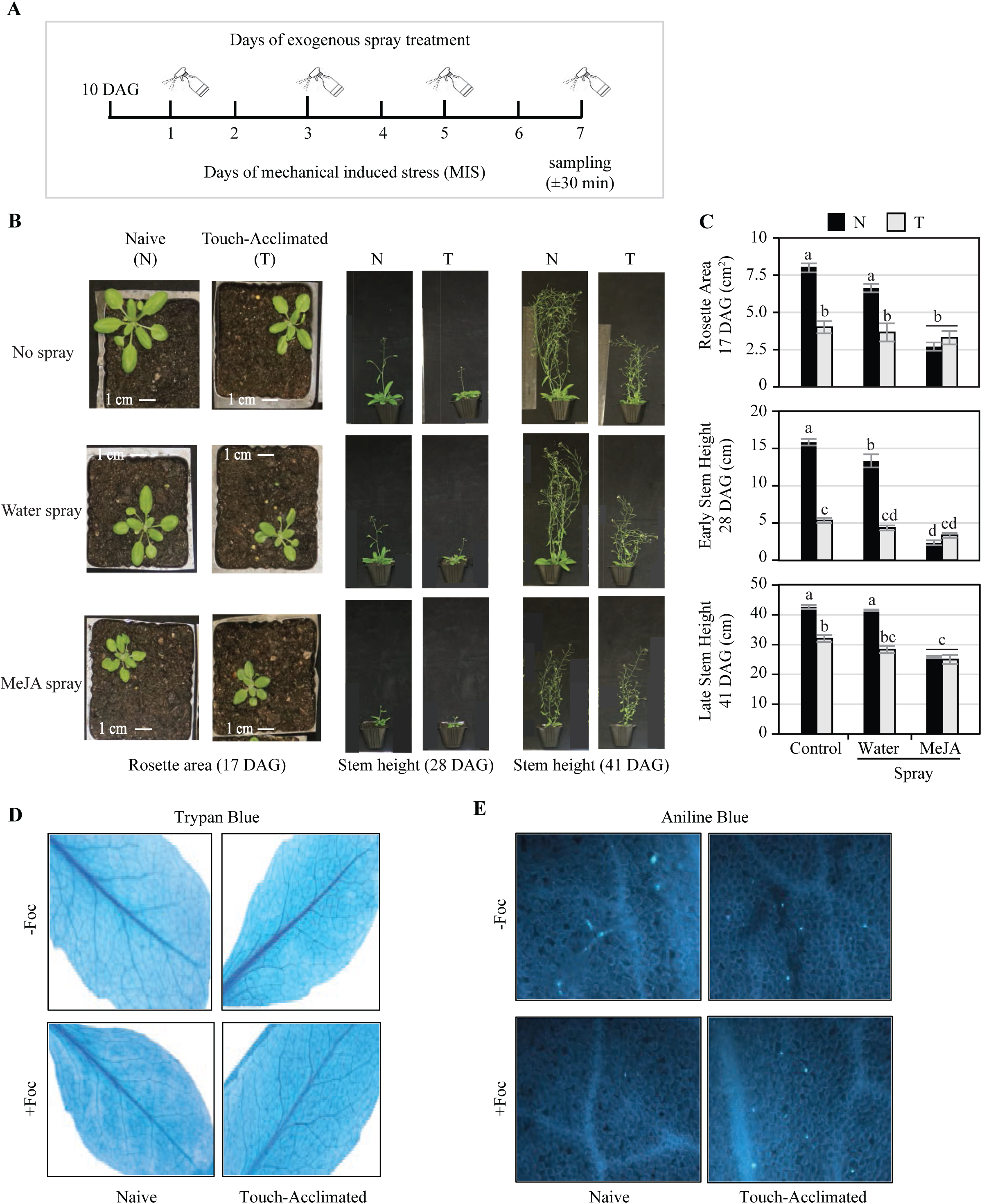
Quantification of plant phenotypes and disease markers in naïve, touch-acclimated and MeJA treated plants. A) Experimental representation showing how naïve juvenile plants (10 days after germination; DAG) are exposed to 7 days of prolonged mechanical induced stress (MIS) and/or sprayed every other day with an exogenous application of a phytohormone or water. B) Representative images of rosette area (17 DAG) and inflorescence stem height (28 and 41 DAG) from unstressed (naïve; N) and touch-acclimated (T) plants sprayed with/without exogenous methyl jasmonate (MeJA) or water. (C) Quantification of Arabidopsis rosette area (cm^2^) and inflorescence stem height (cm) during post-recovery phase at 28 DAG and 41 DAG. D-E) Lactophenol trypan blue (D; stains for necrotic lesions) and aniline blue (E; detects callose accumulation) staining of untreated control and touch-acclimated leaves infected with *Fusarium oxysporum* f. sp. *conglutinans* (Foc) (7-dpi). Error bars show the standard error of the mean (n = 15 plants). Lettering denotes significant differences between treatments by ANOVA (*p* < 0.05). Data is representative of multiple independent experiments.

## SUPPLEMENTARY TABLES

**Supplementary Table 1.**
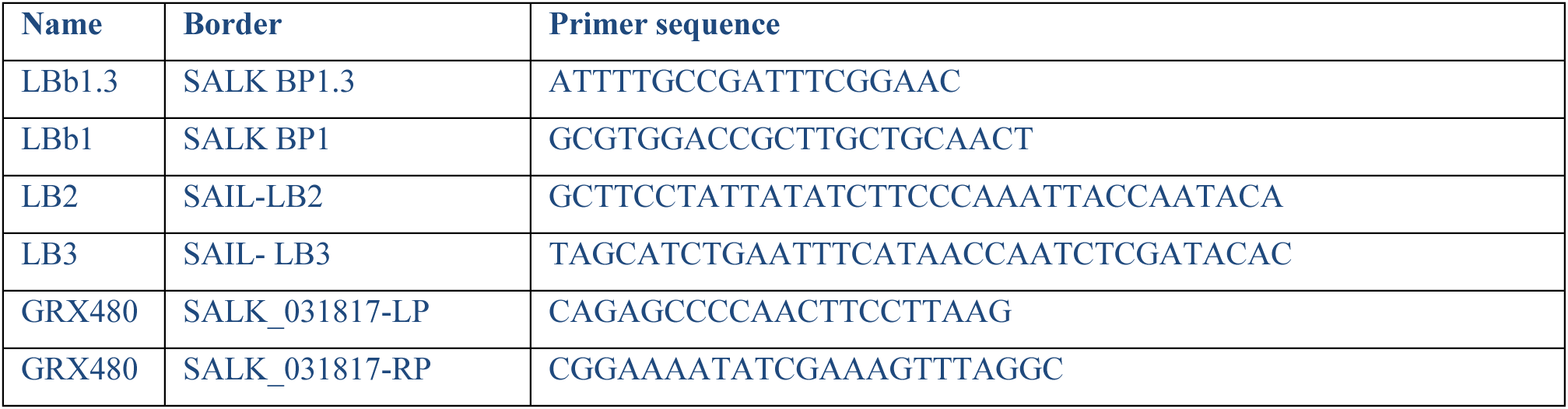
List of primers used to verify T-DNA insertion lines (SALK and SAIL).

**Supplementary Table 2.**
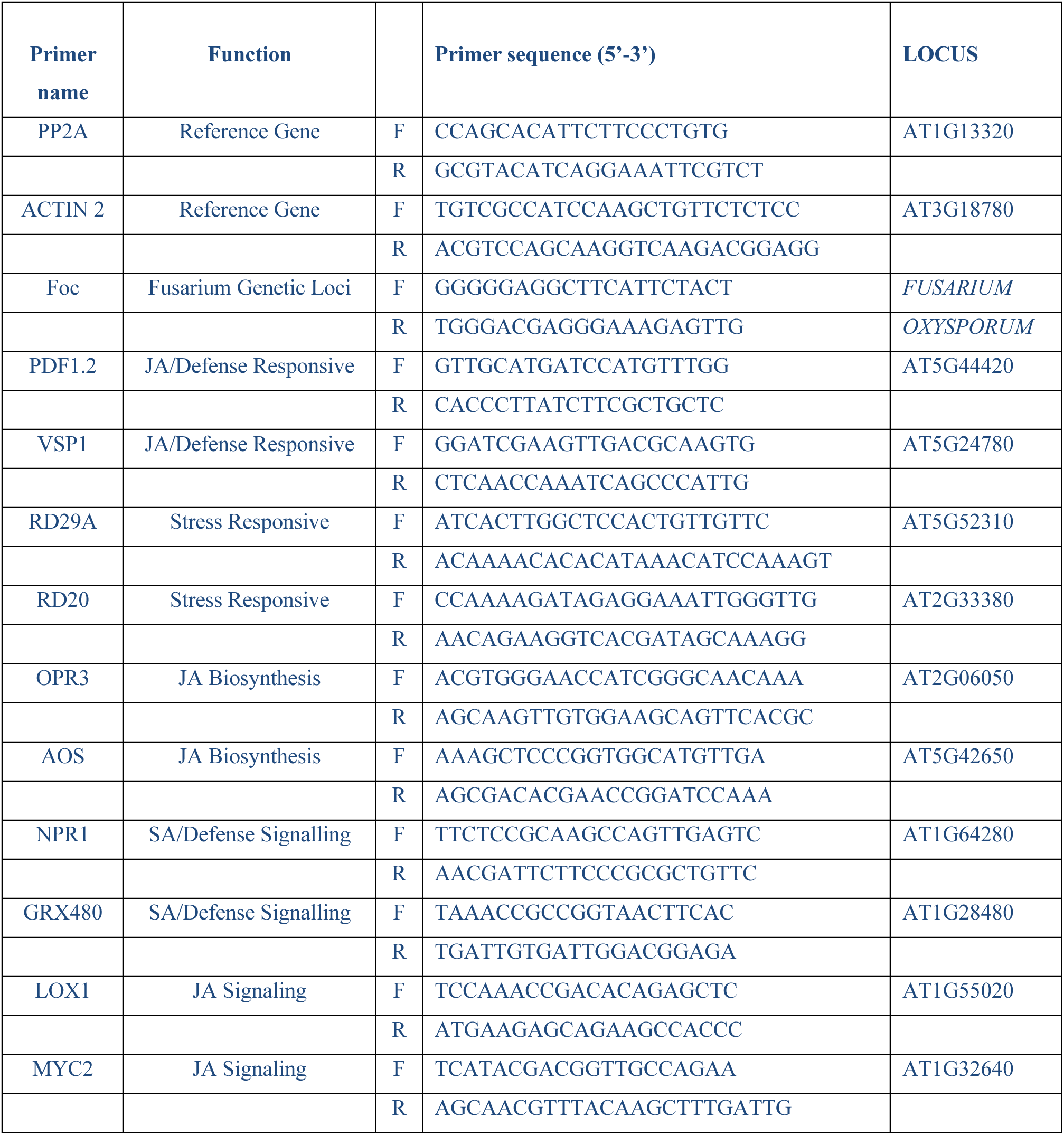
List of primers used for PCR or qPCR of gene expression.

